# PamgeneAnalyzeR open and reproducible pipeline for kinase profiling

**DOI:** 10.1101/589838

**Authors:** Amel Bekkar, Anita Nasrallah, Nicolas Guex, Lluis Fajas, Ioannis Xenarios, Isabel C. Lopez-Mejia

## Abstract

Protein phosphorylation - catalyzed by protein kinases - is the most common post-translational modification. It increases the functional diversity of the proteome and influences various aspects of normal physiology and can be altered in disease states. High throughput profiling of kinases is becoming an essential experimental approach to investigate their activity and this can be achieved using technologies such as PamChip^®^ arrays provided by PamGene for kinase activity measurement. Here we present “pamgeneAnalyzeR”, an R package developed as an alternative to the manual steps necessary to extract the data from PamChip^®^ peptide microarrays images in a reproducible and robust manner. The extracted data can be directly used for downstream analysis.

**Availability and implementation:** PamgeneAnalyzeR is implemented in R and can be obtained from https://github.com/amelbek/pamgeneAnalyzeR.

## Introduction

Analytical pipelines are at the heart of several research endeavors, improving and providing transparent mechanisms for the analysis of complex data. One of the challenges we are facing in today technology providers is the increased necessity for a direct transfer from raw data to an analytical framework. This challenge has been taken on the technology called PamGene that enables the parallel analysis of over 500 kinases and their substrates^1^.

Protein phosphorylation, which consists of a reversible transfer of a phosphoryl group onto specific amino acids on target proteins, is the most common post-translational modification. This mechanism increases the functional diversity of the proteome and influences various aspects of normal physiology by activating and deactivating enzymes, receptors and regulatory proteins. Enzymes known as protein kinases catalyze these events. Protein phosphorylation is often altered in pathological conditions, like cancer, metabolic diseases and neurodegenerative diseases^2, 3, 4^. Kinases phosphorylate their specific targets on specific serine, threonine, and tyrosine residues. Protein kinases can function as receptors on the cell surface or as intracellular mediators. They are essential players in signal transduction and participate in the regulation of all cellular functions, such as proliferation, metabolism, survival and apoptosis.

Unlike methods such as transcriptomics, which only assess the transcribed quantity of each kinase, PamChip^®^ arrays allow a direct examination of kinases at an activity level (kinome profiling). High throughput kinome profiling is becoming an essential experimental approach to investigate aspects of cell and tissue function in physiologic and pathologic conditions. Several techniques, known as phosphoproteomics, have been developed as adaptations of traditional proteomics, to survey protein phosphorylation on a large scale^5,6^. However, phosphoproteomics only measure steady state protein phosphorylation and do not directly reflect cellular dynamics. Alternative approaches using peptide arrays allow the measurement of direct kinase function. Several peptide arrays were developed to measure kinase activity^1,7^. One of those technologies, developed by PamGene, provides PamChip^®^ arrays for kinase activity measurement. PamChip^®^ arrays contain peptides with one or several phosphorylatable residues. Upon incubation with a biological sample this technology enables to measure kinase activity using phosphorylated peptides as readout.

## PamGene PamChip^®^ arrays

PamChip arrays consists of 144 distinct peptides, each composed of 12-15 amino acids, with one or more phospho-sites. PamGene proposes two different types of arrays with peptides containing tyrosine or serine/threonine phosphorylation sites. The PamChips were designed to capture the activity of upstream kinases of either the tyrosine kinome (protein tyrosine kinase or PTK) or the serine/threonine kinome (serine/threonine kinase or STK) respectively. Peptides are printed onto the array in ‘spots’, and like for other microarray technologies, FITC-conjugated antibodies are used to quantify the phosphorylation signal by quantifying the image pixel brightness at each spot. To capture the kinetic of the reactions, the sample fluid is pumped during several cycles. Images are taken at fixed cycle intervals over the course of a reaction. Depending on the chosen experimental design images can be taken at a fix or a varying camera exposure time (10ms, 20ms, 50ms, 100ms, 200ms).

PamGene experiments can generate substantial amounts of data in the form of images taken at different cycles and camera exposure times. Those images have to be explored carefully with the aim of removing and identifying failed chips, which is tedious and prone to operator variation. To enable an operator independent reproducible analysis, we propose “pamgeneAnalyzeR", an open-source R package that extracts the raw data signal from the PamChip images. The extracted data quantification can be used subsequently for downstream analyses such as kinetic exploration or differentially phosphorylated peptides detection. The pamgeneAnalyzeR is an alternative approach to BioNavigator (licensed by PamGene) that aims to automate the data analysis.

## Methodology

The proposed R package is composed of a set of functions that are used to extract data from the images and prepare it for further analysis (figure 1).

**Figure 1:**
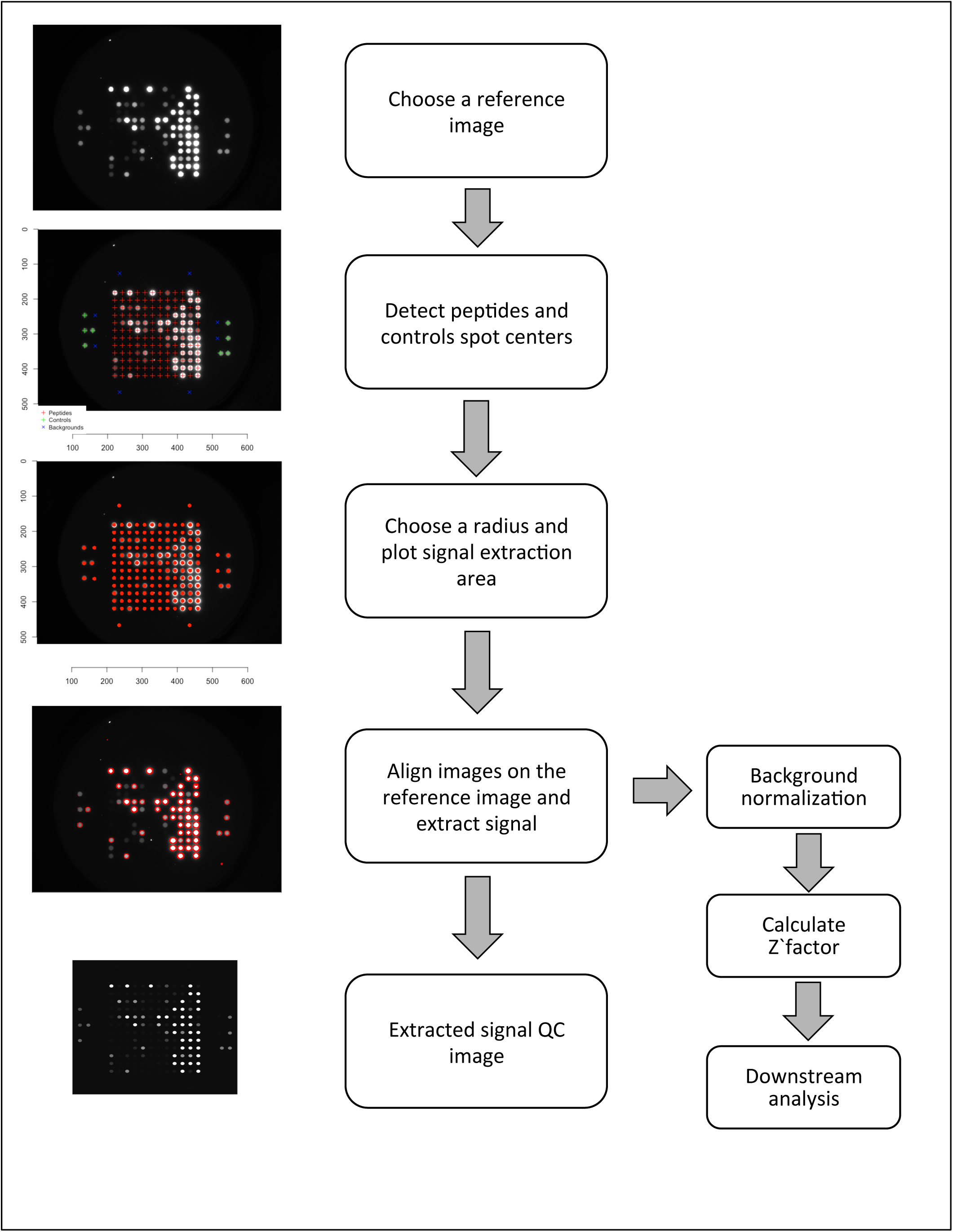
General package workflow. From raw images to extracted signal.

Raw images acquired by the camera are directly used as input of this package. One reference image is selected to detect the centers of peptides spots as well as the right and left controls. The reference image should have at least clear bright left and right control spots, which are used to identify the optimal parameter for the radius of the spot during quantification. This parameter is necessary, since bleaching and spreading of the signal can occur depending on the acquisition protocol. At this step, QC images showing the radius of the spots overlaid onto raw images can be generated. The centers of the spots are used thereafter with the chosen radius to draw a circle on each spot, inside which the pixel brightness values are captured. The pipeline then automatically aligns all the other images onto the chosen reference image and the signal of each spot of each image is extracted. As an assessment of the quality of the process, a synthetic image can be reconstructed from the extracted signal an compared to the original one to ensure that the pixel values where extracted correctly and no drift or shift is observable. Summary statistics of pixel brightness (mean, median, standard deviation and sum) are collected for each peptide spot, the left and right control point as well as 8 control background points.

All collected signals are then merged in a unique R data frame. By default, the median pixel brightness value of each spot is used since it represents a robust measure of the spot signal once aligned to the reference. For background normalization we subtract from each spot the mean of background control spots. A Z’-factor is calculated for each sample representing the dynamic range between positive and negative controls contained by each PamGene chip. Z’-factor values between 0.5 and 1 are considered excellent, values between 0 and 0.5 may be acceptable, and values less than 0 indicate the assay is unlikely to be usable, since positive and negative control readout overlap heavily, indicating that the chip for a given experiment has had some failure.

## Conclusion

Kinome profiling to investigate biological processes is becoming easily accessible thanks to methods such as PamGene PamChip^®^ arrays that provide a valuable tool for direct exploration of cellular kinase activities^8,9,10^. The ability to develop bioinformatic tools to extract and analyze the data in a reproducible and robust manner is critical. We have developed “pamgeneAnalyzeR”, an automated R package as an alternative to the manual steps commonly used to analyze such arrays, with the idea to implement this technique in pipelines applied to cancer and diabetes research. The extracted data can be directly used for downstream analysis. Other tools are also developed allowing downstream analysis of PamChip data, such as the Kinomics toolbox^11^, indicating the interest for this technology and the need for analysis frameworks. The aim of providing this software as an open-source GPL license is to help researchers take full advantage of the technology and also integrate it more in their complex data acquisition environment.

## Acknowledgments

We are thankful to Julien Dorier for his advice.

